# Adult-born granule cells improve stimulus encoding and discrimination in the dentate gyrus

**DOI:** 10.1101/2022.05.12.491644

**Authors:** Diego M. Arribas, Antonia Marin-Burgin, Luis G. Morelli

## Abstract

Heterogeneity plays an important role in diversifying neural responses to support brain function. Adult neurogenesis provides the dentate gyrus with a heterogeneous population of granule cells (GCs) that were born and developed their properties at different times. Immature GCs have distinct intrinsic and synaptic properties than mature GCs and are needed for correct encoding and discrimination in spatial tasks. How immature GCs enhance the encoding of information to support these functions is not well understood. Here, we record the responses to fluctuating current injections of GCs of different ages to study how they encode stimuli. Immature GCs produce unreliable responses compared to mature GCs, exhibiting imprecise spike timings across repeated stimulation. We use a statistical model to describe the stimulus-response transformation performed by GCs of different ages. We fit this model to the data and obtain parameters that capture GCs encoding properties. Parameter values from this fit reflect the maturational differences of the population and indicate that immature GCs perform a differential encoding of stimuli. To study how this age heterogeneity influences encoding by a population, we perform stimulus decoding using populations that contain GCs of different ages. We find that, despite their individual unreliability, immature GCs enhance the fidelity of the signal encoded by the population and improve the discrimination of similar time dependent stimuli. Thus, the observed heterogeneity confers the population with enhanced encoding capabilities.

## Introduction

Cell diversity is ubiquitous in the brain and many studies have highlighted its importance for the encoding of stimuli by a population of neurons [1, 2, 3, 4, 5, 6, 7, 8]. In the dentate gyrus of the hippocampus, cell diversity is enhanced and structured by adult neurogenesis, a mechanism by which new neurons termed granule cells (GCs) are born beyond development [9]. These adult-born GCs mature in a stereotyped way, making a distinctive contribution to hippocampal function during maturation [10, 11, 12, 13]. After about 8 weeks, they become electrophysiologically indistinguishable from other mature neurons [14, 15].

About 4 weeks after cell birth, immature GCs establish functional synapses that can activate postsynaptic targets in CA3 [16, 17, 18]. At this age, they also receive presynaptic inputs from within the hippocampus [19], the septum [19] and the entorhinal cortex [9, 15, 19, 20, 21]. Concurrently, the electrophysiological properties of immature GCs evolve continuously. Experiments in hippocampal slices have shown that 4 weeks old GCs exhibit different electrophysiological properties from those of mature GCs, such as an increased excitation/inhibition balance, a higher membrane excitability, a slower membrane time constant and lower action potential threshold [15, 21, 18, 22]. Therefore, around 4 weeks after cell birth immature GCs become integrated into the hippocampal circuitry and could play a distinctive role in the encoding and transmission of information.

In vivo, immature GCs exhibit distinctive patterns of activity: they fire at higher rates, they are less spatially tuned and are preferentially activated during spatial memory tasks [10, 23]. Immature GCs also promote pattern separation, a computation usually associated with the dentate gyrus which presumably involves augmenting the differences between similar incoming activity before relaying it [24]. Behavioral experiments have shown that ablating or inhibiting immature GCs impairs pattern separation [10, 11, 25], while enhancing neurogenesis or inhibiting mature GCs outputs improves it [12, 13]. Still, the mechanisms by which immature GCs distinct properties shape these functional roles remain largely unexplored.

Unlike random cell diversity, neurogenesis introduces a stereotyped form of diversity that could be leveraged by the dentate gyrus network. Given their distinct properties, immature GCs could perform a differential encoding of incoming activity, promoting pattern separation and other functions. However, experimental studies that address questions related to coding are scarce for immature GCs, partly because recording from immature GCs in vivo is challenging. Pattern separation involves encoding two overlapping stimuli into dissimilar representations. Do mature and immature GCs encode stimuli differently? Furthermore, how do GCs of different age encode a stimulus into a spiking response? The diversity contributed by immature GCs could improve these representations, facilitating their discrimination.

Here, we record the membrane potential of individual GCs while they process fluctuating input currents in hippocampal slices. We use transgenic mice to label immature GCs of different ages and distinguish them from the population. First, we generate a single current stimulus template using a stochastic process with short temporal correlations. Then, we inject the same stimulus template multiple times into each GC to study the structure of the responses across time and trials. To investigate the encoding properties of GCs, we characterize the functional relationship between stimulus and response fitting a statistical model for each recorded GC. Using this encoding model to decode experimental and simulated responses, we study how well these responses preserve stimuli information content. Building in silico populations of GCs of homogeneous and heterogeneous ages, we study the impact of age diversity on stimulus reconstruction and encoding. Finally, we use these populations in a pattern separation task to probe the contribution of immature GCs to the discrimination of time-varying stimuli.

## Results

### Immature GCs responses are less reliable and less aligned with stimulation

Electrophysiological properties of immature and mature GCs are known to differ [15, 26, 25]. Thus, we wondered whether neurons of different age would produce responses with a different temporal structure to the same stimulus. To investigate this, we performed ex vivo whole cell recordings in mature GCs (mGCs, n=21) and adult-born GCs that were 4 and 5 weeks old (4wGCs, n=22 and 5wGCs, n=20) (Figure 1A). Adult-born GCs of different ages were labeled using a Ascl1-CreERT2-Tom mice line (Figure 1B, Methods) [26]. Brain slices were prepared 4 or 5 weeks after tamoxifen injection to obtain tomato expressing GCs of these ages. Excitatory and inhibitory synaptic transmission were blocked pharmacologically with kynurenic acid and picrotoxin, so that differences in recorded activity were only due to differences in the cells intrinsic properties. We measured the passive input resistance and time constant of the GCs by injecting small hyperpolarizing current steps at -70 mV. Both the input resistance and the time constant measured in this way decreased with age (Figure 1-figure supplement 1), consistent with previous reports [15, 26]. Additionally, when injected with depolarizing current steps, immature GCs fired at higher frequencies than mature ones for the same step amplitude (Figure 1-figure supplement 1).

**Figure 1:**
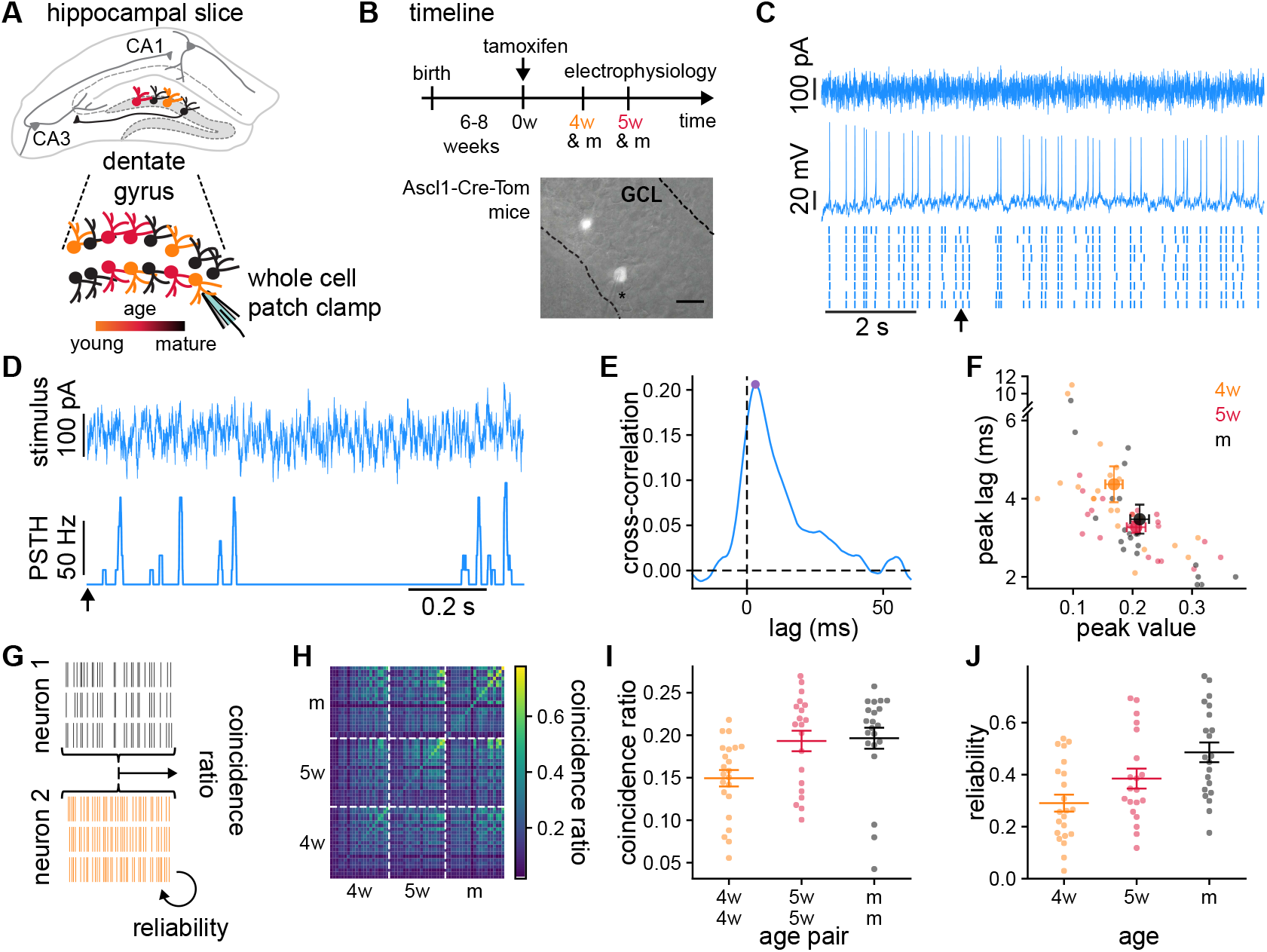
GCs recordings and analysis of the temporal structure of the responses to the same stimulus. **(A)** Schematics of the experimental setup showing a hippocampal slice with the dentate gyrus highlighted in grey, and a blow-up of the dentate gyrus GCs. Colors indicate GCs age: 4wGCs (orange), 5wGCs (red) and mGCs (black). **(B)** Top: experimental timeline. Tamoxifen is injected 6-8 weeks after mouse birth and slices are obtained 4 and 5 weeks after injection. Bottom: hippocampal slice showing labeled immature GCs (bright spots) in the granule cell layer (GCL). Asterisk marks the electrode. Scale bar: 20*µ*m. **(C)** Recording of a mGC. Top: fluctuating current stimulation. Middle: recorded membrane potential from a single trial. Bottom: spike raster plot showing nine trials obtained with the same stimulus. Arrowhead marks the starting time for panel **(D). (D)** Stimulus (top) and resulting PSTH (bottom) from **(C). (E)** Cross-correlation between the stimulus and the PSTH of the mGC in **(C, D)**. Cross-correlation peak (dot) is characterized by its peak value and lag. **(F)** Cross-correlation lag and peak value for all GCs recorded (small dots) and age group averages (large dots). Bars indicate mean ± 1 s.e.m.. Spearman’s correlation: *ρ* = −0.30, *p* = 0.010 between age and lag and *ρ* = 0.25, *p* = 0.047 between age and peak value. **(G)** The average fraction of coincident spikes defines (i) a reliability between different trials from a single GC and (ii) a coincidence ratio between all trials from different GCs. **(H)** Coincidence ratio matrix for all pairs of GCs. The diagonal is the reliability. **(I)** Average coincidence ratio between each GC and all other GCs of the same age (dots). Spearman’s correlation with age of the pair: *ρ* = 0.42, *p* = 7.1 × 10^−4^. **(J)** Reliability of individual GCs (dots). Spearman’s correlation with age: *ρ* = 0.44, *p* = 2.7 × 10^−4^. In **(I, J)** bars indicate mean and ± 1 s.e.m. for each age group.

Next, we used fluctuating stimuli that produce responses with a rich and reproducible temporal structure [27]. We generated a single stimulus template from an Ornstein-Uhlenbeck process with a short correlation time constant (Figure 1C, Methods). Mature GCs have smaller input resistances and integrate larger excitatory and inhibitory currents than immature GCs (Figure 1-figure supplement 1) [15, 21]. Therefore, to make the firing rate of the responses comparable across GCs of different ages, we used the same stimulus template for all GCs while adapting its baseline and amplitude (Figure 1-figure supplement 2). We injected 9 trials of the template stimulus in each GC and recorded the resulting membrane potentials. From these, we extracted the spike timings that we used to study the responses.

We first studied the alignment between the neural responses and the injected stimulus. For each GC, we smoothed the spike trains with a rectangular sliding window and averaged over trials to obtain a Peri-Stimulus Time Histogram (PSTH) (Figure 1D). We then computed the Pearson cross-correlation between the injected stimulus and the PSTHs for different time lags (Figure 1E). The lag of the cross-correlation peak reflects the time scale of stimulus integration, and the peak value the degree of alignment between stimulus and response. The lag and the value at the peak of the cross-correlation were negatively and positively correlated with age respectively (Figure 1F). These results indicate that GCs responses become faster with maturation and exhibit stronger alignment with the stimulus.

While spike times were often preceded by stimulus upswings, the previous analysis doesn’t explore the variability in the responses across trials and different GCs. To investigate this, we introduced a coincidence ratio between pairs of recordings, each recording consisting of all trials (Figure 1G, Methods) [28, 29]. We defined the coincidence ratio as the average proportion of spike coincidences between all possible pairs of trials. A coincidence ratio of 0 means no coincidences between the recordings and a coincidence ratio of 1 means all spikes match their timing. We quantified the reliability in the response of single GCs by computing coincidences between pairs of different trials of the same GC.

The coincidence ratio spanned a wide range of values (Figure 1H). For each GC, we computed its average coincidence ratio with all other GCs of the same age (Figure 1I). Older pairs of GCs exhibited larger coincidence ratios, indicating their responses were more similar to each other. This could be a consequence of individual immature GCs producing less reproducible responses, which can be quantified by the reliability. GC reliability, determined by the diagonal of Figure 1H, took generally higher values than the coincidence ratio as trials from a single GC were usually more similar to each other than to trials from a different GC. Reliability increased with maturation, taking values 0.29 ± 0.03 (mean ± 1 s.e.m.) for 4wGCs, 0.38 ± 0.04 for 5wGCs and 0.49 ±0.04 for mGCs (Figure 1J). Consistent with the cross-correlation analysis, this indicates that GCs are able to produce more robust responses with maturation. This effect could not be attributed to differences in firing rate, as shown by the reliabilities obtained after randomizing the timing of the spikes in every trial (Figure 1-figure supplement 2). Thus, our data indicates that immature GCs are less aligned with the stimulus and produce less reliable responses than mature ones.

### Immature GCs integrate over longer time scales and exhibit weaker refractory effects

The observed differences in the stimulus alignment and reliability of the responses suggest that GCs of different ages do not perform the same stimulus encoding. Thus, we sought to find a functional relationship between the stimulus and the GCs responses to characterize their encoding properties, by fitting a spike response model (SRM) to our data (Figure 2) [30]. The SRM is a statistical model in the family of Generalized Linear Models (GLMs) which allow for a direct fitting procedure and have been widely used to provide accurate statistical descriptions of spiking data [8, 31]. A key feature of these models is that they provide a quantitative characterization of neural activity in terms of parameters that can be linked to biophysical properties.

**Figure 2:**
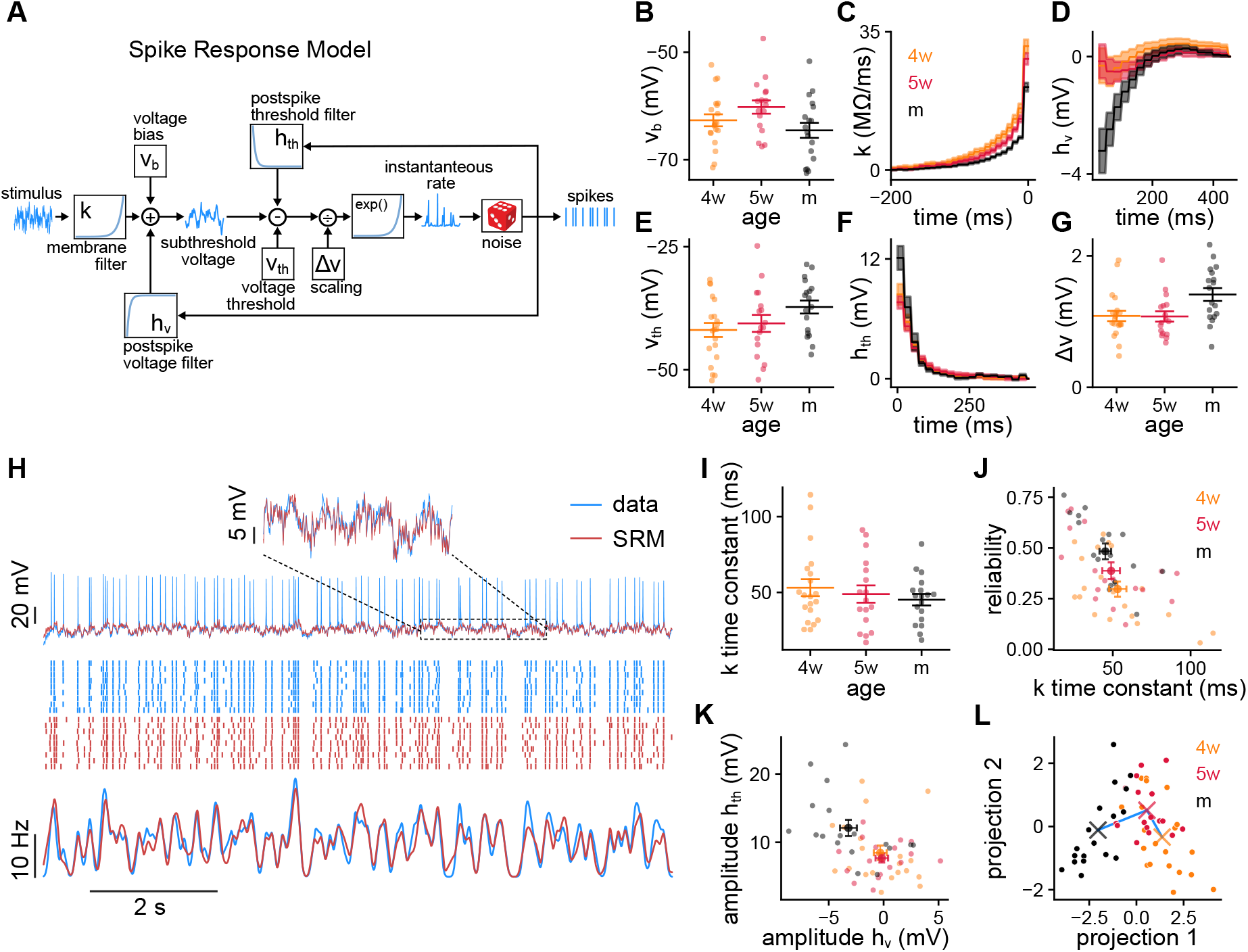
Spike response model fits to recorded GCs. **(A)** Schematics of the spike response model (SRM). **(B-G)** SRM parameters obtained for all GCs of different ages: **(B)** voltage bias *v*_*b*_, **(C)** membrane filter *k*(*t*), **(D)** postspike membrane potential deflection *h*_*v*_(*t*), **(E)** static voltage threshold *v*_*th*_, **(F)** postspike threshold deflection *h*_*th*_(*t*) and **(G)** voltage scaling factor Δ*v*. **(H)** Validation data (blue) and SRM prediction (red). Top: Subthreshold membrane potential. Middle: Spike raster plots of the recorded responses and SRM simulations. Bottom: PSTHs of the spike trains. **(I)** Time constants extracted from the filter *k*(*t*). Spearman’s correlation between age and population time constant (bootstrapped): *ρ* = −0.567, *p* = 6.3 × 10^−2^. **(J)** GC reliability vs. time constant of the filter *k*(*t*). Spearman’s correlation: *ρ* = −0.61, *p* = 6.4 × 10^−7^. **(K)** Amplitudes of the filters *h*_*v*_(*t*) and *h*_*th*_(*t*). Spearman’s correlation: *ρ* = −0.37, *p* = 6.0 × 10^−3^ between age and 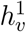, and *ρ* = 0.35, *p* = 8.1 × 10^−3^ between age and 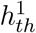. (**L**) GCs parameters projected on the Linear Discriminant Analysis components subspace. The crosses represent the means of the age groups, connected by a blue line. In **(B, E, G, I-L)** small dots represent single neurons. In **(B, E, G, I)** long bars indicate means and short bars means ± 1 s.e.m.. In **(J, K)** large dots and bars indicate means ± 1 s.e.m.. In **(C, D, F)**, lines with shaded areas indicate means ± 1 s.e.m..

The SRM describes both the subthreshold membrane potential and the spiking response to a current stimulation (Figure 2A, Methods). The stimulus first passes through a membrane filter *k*(*t*) that defines how the input current is dynamically transduced in voltage variations. Next, the subthreshold membrane potential is generated by summing the filtered stimulus, a constant bias *v*_*b*_ and the postspike voltage deflection *h*_*v*_(*t*). The bias is a baseline voltage and the postspike voltage deflection accounts for the effect of intrinsic currents occurring after a spike. A moving voltage threshold is then subtracted from the subthreshold membrane potential. The exponential of this difference scaled by a factor Δ*v* sets the time dependent spiking rate. The moving threshold is the sum of a constant threshold *v*_*th*_ and the postspike threshold deflection *h*_*th*_(*t*) that accounts for threshold refractory effects. Finally, spikes are randomly generated at every discrete time point from the time dependent spiking rate.

We fitted the SRM to single trial 99 s recordings from individual 4wGCs (n=20), 5wGCs (n=17) and mGCs (n=18), using rapidly fluctuating stimuli generated as described in the previous section. We performed the SRM fitting in two steps: first we extracted the subthreshold parameters *v*_*b*_, *k*(*t*) and *h*_*v*_(*t*) using the subthreshold membrane potential (Figure 2B-D, Methods); then we extracted *v*_*th*_, *h*_*th*_(*t*) and Δ*v* by maximizing the likelihood of the spike times while keeping the subthreshold parameters fixed (Figure 2E-G, Methods). The SRM yields predictions on both the subthreshold membrane potential and the spikes of the response. To validate the fits, we used the 9 trials 10 s recordings described in the previous section. The SRM captured both the subthreshold membrane potential and spiking responses of the validation data (Figure 2H). We used the validation data to compute the root mean squared error of the subthreshold membrane potential prediction and a normalized log-likelihood (Figure 2-figure supplement 1, Methods). To assess the quality of the spike trains generated by the SRM, we also computed the coincidence index *M*_*d*_ between simulated and recorded spike trains [29, 30]. The reliability of the simulated spike trains was generally smaller than the reliability of the data but they were linearly correlated, indicating that the differences between GCs were mostly preserved (Figure 2-figure supplement 1).

The parameters obtained from the fit reflect differences in the way that GCs of different ages encode stimuli. While passive properties determine the membrane potential deflection for small current steps in the absence of spikes, the membrane filter *k*(*t*) determines how the stimulation current is actively transduced into voltage during spiking activity. *k*(*t*) is composed of a rapid decay attributed to the patch pipette [32] and a tail that is well approximated by an exponential decay (Figure 2C). The time constant of the slower exponential decay measures the time scale of stimulus integration. The area under the exponential is an electrical resistance and determines the average membrane potential deflection in the absence of spikes. Both the population time constant and the resistance of the filter *k*(*t*) decreased with age (Figure 2I, Figure 2-figure supplement 1), consistent with the passive properties observations (Figure 1-figure supplement 1) [15, 26]. These properties were correlated with their passive counterparts but were generally smaller (Figure 2-figure supplement 1). Slower membranes could partly explain the observed unreliability of immature GCs, as neurons with longer time constants tend to be less reliable (Figure 2J). The filters *h*_*v*_(*t*) and *h*_*th*_(*t*) quantify the postspike deflections in membrane potential and spiking threshold respectively. We quantified their amplitudes by using the first values of the filters (Figure 2K). While mGCs membranes tended to hyperpolarize after a spike, this effect was generally smaller for immature GCs (Figure 2D,K). This hyperpolarization could be attributed to spike triggered potassium currents, which have been reported to be smaller in immature GCs [15, 26]. While the threshold after a spike increased for all GCs, the amplitude of the filter *h*_*th*_(*t*) was also positively correlated with age.

Next, we wanted to explore whether the obtained parameters could be used to discriminate GCs ages. We used *v*_*b*_, *v*_*th*_, Δ*v* and the first coefficients of the filters *k*(*t*), *h*_*v*_(*t*) and *h*_*th*_(*t*) to classify GCs ages using a Linear Discriminant Analysis (LDA) (Figure 2L) [33]. SRM parameters could be used to classify mGCs and 4wGCs with cross validated accuracies of 0.72 and 0.75 respectively. The other 0.28 and 0.25 fractions were classified as 5wGCs indicating that 4wGCs and mGCs were very well discriminated by the LDA. Being maturationally in between, the classification accuracy of 5wGCs was 0.47, with a fraction of 0.41 5wGCs being classified as 4wGCs. This suggests 5wGCs were closer in parameter space to 4wGCs than to mGCs. The coefficients of the filter *k*(*t*) play an important role in the classification as they have large weights in the LDA components (Figure 2-figure supplement 2). The LDA indicates that the SRM parameters reflect maturational differences and can be used to distinguish GCs ages with higher than chance accuracy.

We observed that mGCs have a smaller input resistance, a faster membrane and experience stronger postspike refractory effects. Our results suggest GCs of different ages might transmit different properties of an afferent stimulus. Besides providing us with parameters that can be compared across ages, the SRM provides us with a statistical description of the GCs. This description can be used to generate spike trains and further explore the way in which GCs represent stimuli. Next, we use our GCs models in a decoding framework to explore the fidelity of stimulus encoding and quantify the information content in the spike trains of the different age groups.

### Immature GCs transmit less information per spike and produce less precise reconstructions

As spiking responses encode information about the stimulus, we wondered, What can we infer about the stimulus by observing the responses of GCs of different ages? To investigate this question, we used the SRM to perform Bayesian model-based decoding [8, 31, 34]. Model-based decoding finds the most probable stimulus that produced one or more spike trains, given a prior distribution (Figure 3A, Methods). This decoded stimulus can then be compared with the original to evaluate the fidelity of the reconstruction. Furthermore, the procedure can be used to estimate the mutual information between stimulus and responses [34]. More broadly, we can use this framework to asses the fidelity of the code, that is to determine how well GCs of different ages encode information about the stimulus in the spike trains they produce.

**Figure 3:**
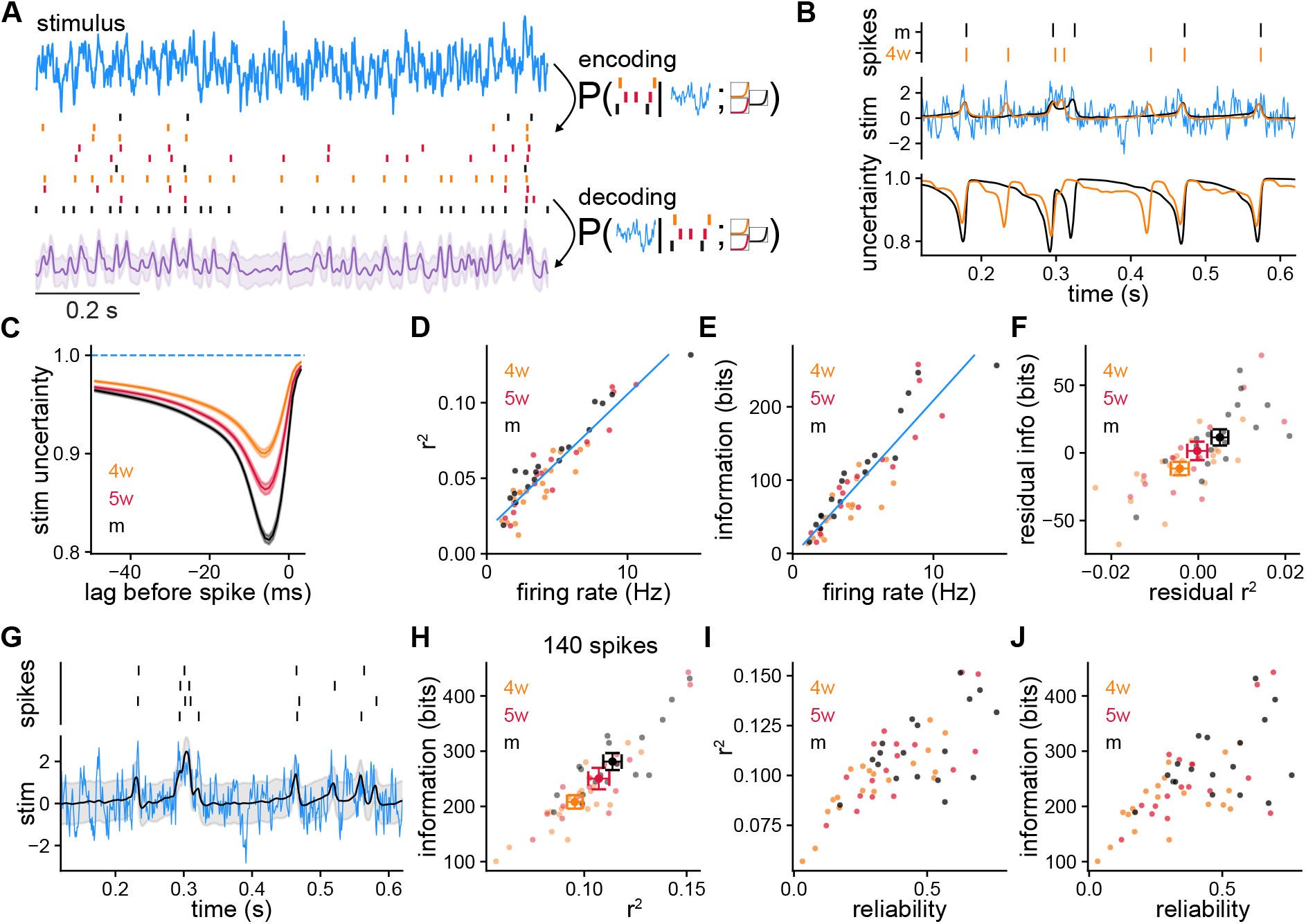
Model-based Bayesian decoding of the stimulus using single GCs. **(A)** Model-based Bayesian decoding diagram. Neurons encode a stimulus in spike trains that can be used to estimate the stimulus that produced them. **(B)** Spike trains (Top) used separately to obtain two stimulus estimations (Middle) and their respective uncertainties (Bottom) about the stimulus. mGC decoding in black, 4wGC decoding in orange, true stimulus in blue. **(C)** Average uncertainty about the stimulus before a spike at lag 0. Spearman’s correlation between minimum uncertainty and age: *ρ* = −0.51, *p* = 6.0 × 10^−5^. **(D)** Coefficient of determination *r*^2^ obtained by decoding with individual spike trains from single GCs vs. firing rate. Pearson’s correlation: *ρ* = 0.93, *p* = 3.6 × 10^−25^. The blue line is the linear fit using all GCs. **(E)** Estimated mutual information between the stimulus and the GCs responses vs. firing rate. Pearson’s correlation: *ρ* = 0.91, *p* = 2.4 × 10^−22^. The blue line is the linear fit using all GCs. **(F)** Residual information vs. residual *r*^2^ after subtracting the linear relationships of **(D, E)**. Spearman’s correlation between age and residual *r*^2^: *ρ* = 0.39, *p* = 2.9 × 10^−3^; Spearman’s correlation between age and residual information: *ρ* = 0.38, *p* = 3.8 × 10^−3^. **(G)** Decoding example using spike trains from a single GC produced by multiple trials of the same stimulus. **(H)** Information vs. *r*^2^ obtained by decoding with 140 spikes on average using a different number of trials for each GC to compensate for firing rate differences. Spearman’s correlation between age and *r*^2^: *ρ* = 0.37, *p* = 6.0 × 10^−3^; Spearman’s correlation between age and information: *ρ* = 0.45, *p* = 6.1 × 10^−4^). *r*^2^ from **(H)** vs. GC reliability. Spearman’s correlation: *ρ* = 0.71, *p* = 1.2 × 10^−9^. **(J)** Information from **(H)** vs. GC reliability. Spearman’s correlation: *ρ* = 0.62, *p* = 5.1 × 10^−7^.

We first performed the decoding using single experimentally recorded spike trains. We found the most probable stimulus that produced the spikes and the uncertainty about its value as a function of time (Figure 3B). During intervals in which a GC is silent, there is no information about the stimulus; hence the estimation takes baseline values and the uncertainty is highest. Just before an isolated spike, the stimulus is usually predicted to be above average and the uncertainty is reduced. Averaging over spikes and GCs, we found that the minimum uncertainty about the stimulus before a spike was reduced more in mature GCs (Figure 3C).

While recorded spike trains allowed us to perform the decoding of the experimental stimulus, the SRM can be used to simulate responses to multiple different stimuli for decoding. Thus, we sampled different stimuli from the same stochastic process that was used experimentally, and repeated the decoding procedure using single spike trains generated with the SRM parameters of each GC. This strategy allows us to quantify decoding performance by estimating the mean coefficient of determination *r*^2^ of the reconstruction and the mutual information between the stimulus and the GC spike trains (Methods).

While the coefficient *r*^2^ involves computing the error between the reconstructed and the original stimuli, the mutual information measures the correspondence between stimulus and response and quantifies the reduction in uncertainty about the stimulus after observing the given spike trains [35]. Both the total *r*^2^ and mutual information strongly correlated with GC firing rate (Figures 3D,E). This is expected as we used only the spikes to decode the stimulus. Decoding performance using multiple stimuli and simulated spike trains was very similar to the one obtained by using the experimentally recorded stimuli and spike trains (Figure 3-figure supplement 1). To study the difference between GCs ages, we removed the average contribution of firing rate to decoding performance. For this, we subtracted the linear relationships of Figures 3D,E obtained with all GCs from the *r*^2^ and information of each GC, obtaining the respective residuals (Figure 3F). Both the residual *r*^2^ and information correlated with GC age indicating that for a given firing rate, responses from mature GCs generally produced more precise reconstructions and were more informative about the stimulus.

We showed that GCs have different degrees of reliability and the same stimulus will elicit different individual responses from the same neuron (Figure 1C). Is it possible to extract more information from a single GC by observing multiple responses to a single stimulus? Does GC reliability influence this? Using responses to multiple trials of the same stimulus improved both the *r*^2^ and the information of the decoding (Figure 3G). For small fold increments in the number of trials used for decoding, the information about the stimulus increased by almost the same fold (Figure 3-figure supplement 2). Immature GCs benefit more from using multiple trials for decoding as a consequence of their noisier responses. To study age differences while equalizing differences in firing rate, we decoded stimuli using for each GC a number of trials such that the total number of spikes was 140, on average (Figure 3H, Methods). Consistent with the results of Figure 3F, when equalizing the number of spikes for each GC, decoding with immature GCs resulted in lower *r*^2^ and information values. When using larger numbers of spikes to equalize the firing rate differences the correlations between age and decoding performance decreased (Figure 3-figure supplement 2). We also found that when equalizing the number of spikes used for decoding, the resulting *r*^2^ and information values were larger in the GCs that produced more reliable responses (Figure 3I,J).

We next studied how the different age groups complement each other by using pairs of different GCs to decode. We used a single stimulus to generate 140 spikes on average from each GC in the pair and performed the decoding (Figure 3-figure supplement 3). Using pairs of GCs of the same age, we found that more mature pairs achieved higher values of *r*^2^ and information. Pairing mGCs with immature GCs tended to degrade decoding performance resulting in smaller *r*^2^ and information values. Coherent with the results from single GCs, more immature pairs achieved overall worse decoding performance.

Our results indicate that immature GCs produce less precise reconstructions and convey less information about the stimulus, suggesting that they produce generally worse stimuli representations. Moreover, mGCs mostly increase decoding performance when paired with other mGCs rather than with immature ones. Thus, these results cast doubts about whether immature GCs can enhance coding in a population.

### Immature GCs improve stimulus reconstruction in a population

Our results so far suggest that mGCs, isolated or in pairs, are generally better than immature GCs for reconstructing stimuli. However, in a larger population there might be synergistic effects, and GCs that did not perform well by themselves could still play a significant role in enhancing population coding. Thus, we wondered whether immature GCs could improve decoding in a population consisting of a larger group of neurons. When building a population, even for a moderate number of neurons, trying all possible combinations between them results in prohibitive computational costs. Hence, we adopted a sequential greedy procedure to construct populations of GCs that optimize stimulus reconstruction [8]. To build a population, we started with a single mGC and sequentially added individual neurons from the full pool of mature and immature GCs in the experiment (Figure 4A, Methods). At each step, using the group built so far, we performed the decoding adding each possible GC to the population, and kept the one that yielded the largest *r*^2^ for the extended group. As decoding performance depends on the number of spikes (Figure 3D), we used a different number of trials for each GC so that the total number of spikes that each one contributed was 1200 on average (Figure 4B). In this way, we reduced the preference for trivially choosing GCs with high firing rates.

**Figure 4:**
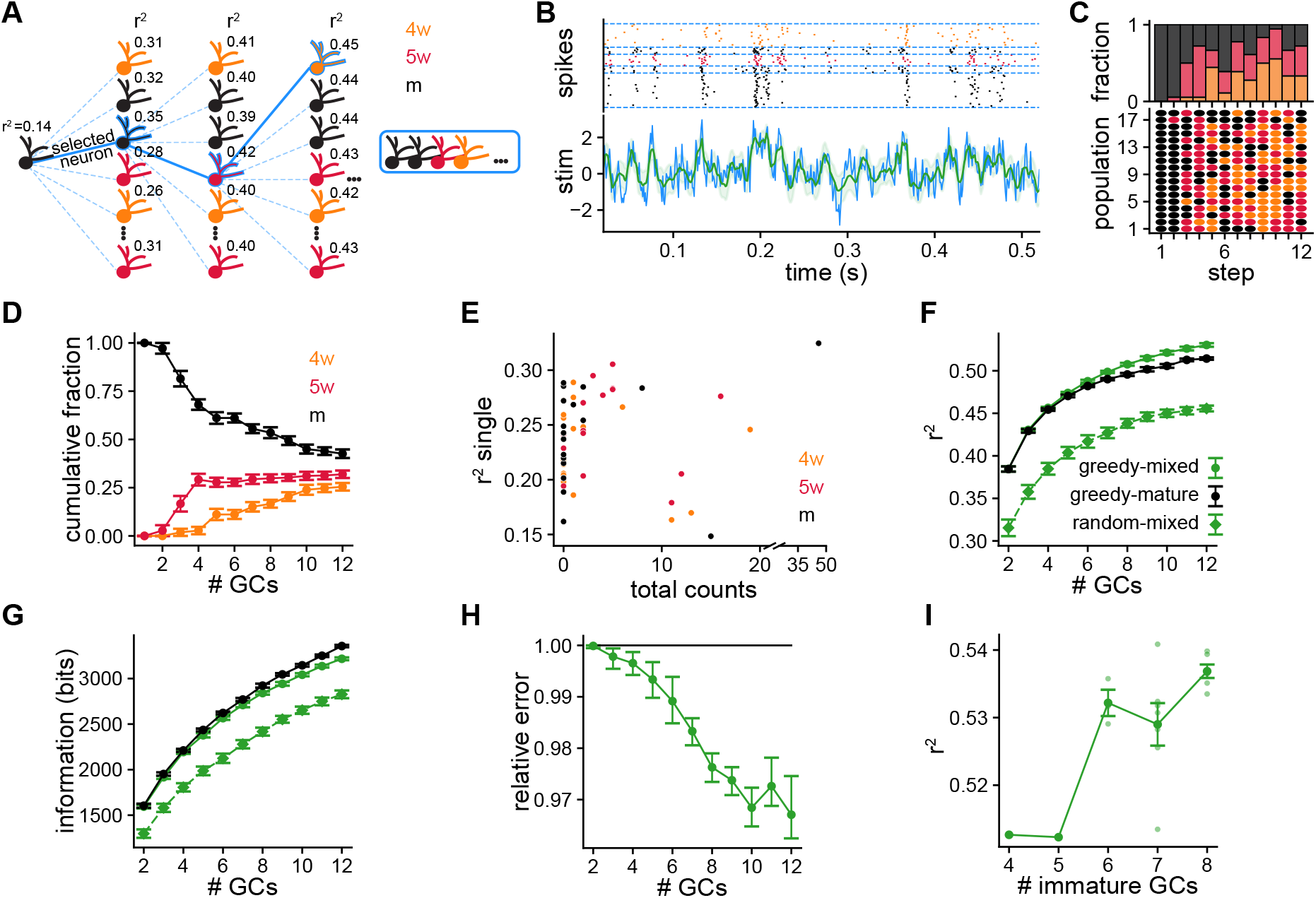
Greedy procedure used to build populations of GCs optimized for stimulus reconstruction. **(A)** Greedy procedure diagram: at each step, the GC that optimizes stimulus reconstruction measured by average *r*^2^ is chosen. **(B)** Decoding example using a population of 5 GCs of different ages. Top: Raster plot showing the spike timings of each GC separated by blue dashed lines. Each GC contributes a different number of trials to equalize the total number of spikes. Bottom: The original stimulus (blue line) is shown together with the reconstructed stimulus (green line) and its uncertainty (shaded green). **(C)** Bottom: Selected GCs at each step. Rows are populations of GCs selected with the greedy procedure. Dot color represents GCs ages. Top: Age fractions of selected GCs at each step. **(D)** Average cumulative fraction for each age for increasing number of GCs in the populations. Error bars indicate 1 ± s.e.m. **(E)** Average *r*^2^ value achieved by single GCs vs. total number of times the GC was selected. **(F, G)** Average over populations of **(F)** *r*^2^ and **(G)** mutual information for increasing number of GCs in the populations. Error bars indicate ±1 s.e.m. **(H)** Reconstruction mean squared error of greedy mixed-age populations relative to greedy mature populations computed by bootstrapping populations from both groups. Error bars indicate symmetric 95% c.i. **(I)** Average *r*^2^ vs. number the of immature GCs for populations of 12 GCs. Spearman’s correlation: *ρ* = 0.67, *p* = 2.3 × 10^−3^. Darker dots indicate averages and error bars indicate ±1 s.e.m..

We built mixed age populations following the greedy procedure. In the first step, we started with the 18 different mGCs we used to fit the SRM (Figure 4C). As expected from the results decoding with pairs of GCs (Figure 3-figure supplement 3), mGCs were chosen almost exclusively among the full pool in the second step. Unexpectedly, from the third step onward, immature GCs were chosen by the greedy procedure with increasing preference (Figure 4C). The fraction of immature GCs in the populations increased at the expense of the mGCs (Figure 4D). For populations of 12 GCs, approximately 43% were mGCs, 32% were 5wGCs and 25% were 4wGCs. We found that 51% of all the GCs were never chosen in any population by the greedy algorithm, 34.5% of the GCs were selected between 1 and 10 times and 14.5% were selected over 10 times (Figure 4-figure supplement 1). This indicates that the final populations strongly differ from random selection, which would have each GC selected 3.6 times on average. Furthermore, these results highlight the diversity of the final populations, since a large fraction of GCs were selected a smaller number of times than the number of populations, hence these are not composed of the same neurons. Moreover, although some of the preferentially chosen GCs achieved large *r*^2^ values when used individually, some highly selected GCs yielded relatively poor reconstructions by themselves (Figure 4E) as well as in pairs (Figure 4-figure supplement 1). These observations indicate that decoding performance is a result of a non-trivial synergy between the GCs composing a population, and optimal performance is not achieved by taking the best individual GCs.

The greedy procedure selected immature GCs to improve stimulus reconstruction. Both *r*^2^ and the mutual information steadily increased with the number of GCs in the populations, and were clearly larger than those of populations of randomly selected GCs (Figures 4F,G). Still, the procedure performs a stepwise optimization, therefore choosing immature over mature GCs early in the construction could result in worse decoding performance for the final populations. We thus compared the decoding performance of the greedy mixed-age populations with exclusively mGCs populations, built following the greedy procedure and starting from the same pool of initial mGCs. With increasing number of neurons, mixed-age populations achieved consistently larger *r*^2^ values in comparison with exclusively mature ones (Figure 4F). This resulted in a steady average decrease in the relative reconstruction error, reaching a final relative improvement of 3% (Figure 4H). Moreover, we found a positive correlation between the *r*^2^ of a population and the number of immature GCs that it contains (Figure 4I). However, the populations of mGCs achieved larger information values than the mixed-age ones (Figure 4G). Thus, while single mGCs perform better than immature ones, mixed-age populations improve stimulus reconstruction, suggesting that the diversity contributed by immature GCs could be beneficial for transmitting distinct properties of stimuli with rich temporal structure.

### Immature GCs in a population improve stimuli discrimination

We showed that the presence of immature GCs in a population can enhance stimulus reconstruction. The dentate gyrus is thought to play a key role in pattern separation [24]. Thus, we wondered whether a downstream region could benefit from reading the output of a mixed-age population to enhance discrimination. We designed a pattern separation task within the decoding framework by using it to discriminate between pairs of correlated stimuli (Figure 5A, Methods). We used mixed-age and mature-only populations of 10 GCs that we constructed in the previous section. We generated pairs of correlated stimuli and then used one of them to generate spike trains. We then performed the decoding to obtain a stimulus reconstruction. To discriminate, we compared the errors between the reconstruction and each of the two stimuli, expecting that this discrimination task would become increasingly harder as the correlation between stimuli increases.

**Figure 5:**
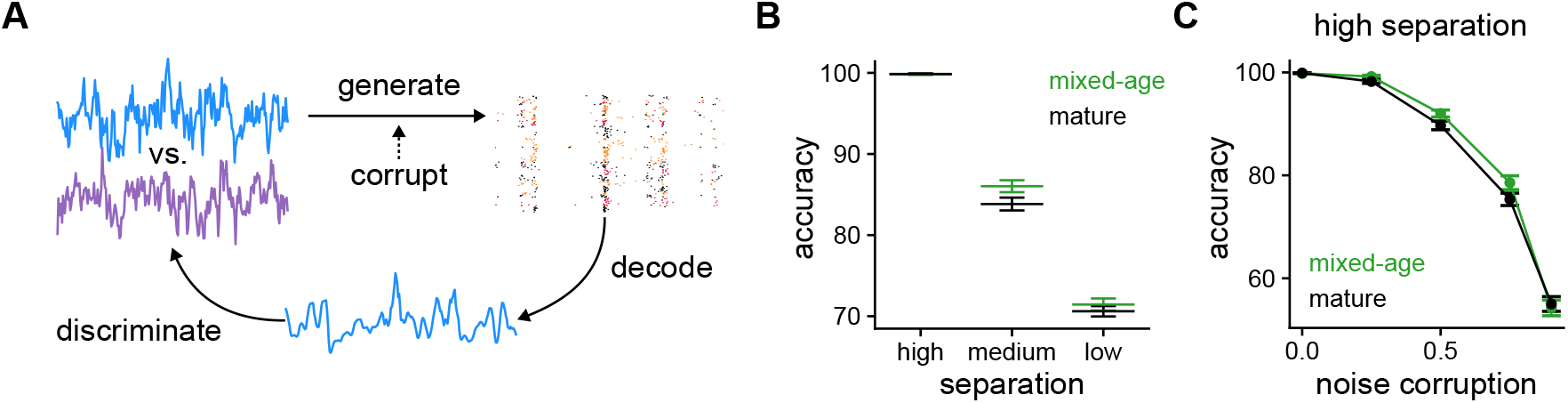
Pattern discrimination between pairs of fluctuating stimuli. **(A)** Diagram of the pattern discrimination procedure. **(B)** Discrimination accuracy achieved by mixed and exclusively mature populations for different degrees of separation between the two stimuli. Mann-Whitney U test high separation: *p* = 0.36, medium: *p* = 1.0×10^−2^ and low: *p* = 0.19. **(C)** Discrimination accuracy vs. level of noise corruption in the stimuli for high degree of separation. Mann-Whitney U test 0.25: noise *p* = 3.6 × 10^−3^, 0.5: *p* = 2.3 × 10^−2^, 0.75: *p* = 6.1 × 10^−2^ and 0.9: *p* = 0.47.

We then explored whether the populations could be used to correctly discriminate between the stimuli with different degrees of separation. We defined three degrees of separation (low, medium and high) that correspond to different correlation values between the jointly generated stimuli (Figure 5B, Methods). For high separation the two stimuli are easily discriminated with almost 100% accuracy both by the mixed-age and mature-only populations. For low separation, stimuli are hard to discriminate and while the obtained accuracy is above chance, the differences between the mixed-age and mature-only populations are small. However, for medium degree of separation the mixed-age populations outperformed the mature-only at the discrimination task, suggesting that age heterogeneity in the dentate gyrus could also be leveraged to separate time-varying stimuli.

Finally, we tested whether immature GCs could also improve discrimination if the stimulus is corrupted with noise. To do this, for each GC we produced a different corrupted version of the stimulus used for generating spike trains. As now each spike train is generated with a corrupted version of the stimulus, correct discrimination will depend on the capability of the decoder to average out the noise in the population. For a fixed degree of separation between the pairs of stimuli, adding noise is detrimental to discrimination performance for the populations (Figure 5C). Notably, the mixed-age populations consistently outperform the mature-only for moderate amounts of noise. In summary, our results suggest that immature GCs in the dentate gyrus introduce a degree of heterogeneity at the population level that can be leveraged to reconstruct an incoming stimulus better and, at the same time, discriminate it from others.

## Discussion

Neurogenesis contributes a substantial number of granule cells to the dentate gyrus [36]. Previous works have studied the electrophysiological properties of adult-born immature GCs in hippocampal slices [15, 21, 22]. Other works have addressed the impact of perturbing neurogenesis on behavior [11, 13]. However, in vivo studies that record from immature GCs are rare [10], and the mechanisms by which neurogenesis influences coding are largely unknown. Here we approach this question combining measurements in hippocampal slices and statistical modeling to study the impact of age heterogeneity on stimulus encoding. We explore the idea that immature GCs could actively aid representation and discrimination of stimuli. We record the responses of dentate gyrus GCs of different ages to stimuli with rich temporal structure. The fluctuating stimuli we use produces reliable responses, allowing us to study their structure over time and across trials. We find that immature GCs produce more variable responses, less reproducible across trials and less aligned with the stimulus. We then fit a spike response model to capture the subthreshold membrane potential and spiking responses. We find that immature GCs integrate stimuli over longer time scales and exhibit weaker refractory effects. We show that these parameters can be used to discriminate GCs ages. Decoding stimuli from the responses of single GCs, we find that stimulus reconstruction and information improve with maturation. Unexpectedly, despite the worse individual performance of immature GCs, we find that they aid decoding in a population. Finally, we design a pattern separation task using our framework and show that immature GCs enhance discrimination of highly correlated stimuli.

To mimic inputs with complex temporal structure, we use fluctuating current injections and build statistical models to investigate the differences between GCs of different ages. We block inhibitory and excitatory transmission to control the stimulus and investigate the interaction between the stimulus and the intrinsic properties of GCs of different ages. We observe that immature GCs produce generally less reliable responses, that are also less correlated with the stimulus and with the responses of other GCs. Immature GCs responses are also more variable in vivo, exhibiting less spatial specificity when animals perform a spatial exploration task [10]. Immature GCs responses are less controlled by inhibitory circuits than mature GCs, spiking in response to weaker stimuli and with more variable timing [21, 22]. Thus, the differences in reliability that we report here could be further increased by the network. Additionally, the amount of excitatory recurrence in the dentate gyrus is small compared to other hippocampal areas like CA3, hence we do not incorporate recurrent connectivity when decoding with populations of GCs [37].

Using the SRM, we quantitatively characterize GCs responses to the applied stimuli. Studying the SRM parameters, we observe that the membrane filters of immature GCs have larger amplitudes and longer time scales, their thresholds are lower and they experience weaker refractory effects. Previous works using current steps have reported that immature GCs have larger input resistances and time constants, lower action potential thresholds and smaller postspike potassium currents [15, 26]. These observations validate our approach and show that parameters obtained from the fit are linked to biophysical properties. Moreover, our SRM characterization reproduces the spiking responses, allowing us to further study the stimulus-response transformation performed by GCs. We find that immature GCs low reliability can be partly attributed to their longer time scales of stimulus integration.

The model-based decoding framework that we use allows us to study how GCs represent stimuli without making precise assumptions about the nature of the neural code. By decoding stimuli with single GCs and populations, we find that reliable neurons generally achieve better decoding performance and that optimal populations combine diversity with homogeneity, using neurons with different properties while often including the same neuron multiple times. Similarly, a previous study reported similar observations using a model-based approach to explore the impact of intrinsic diversity in the olfactory bulb [8]. Our study explores not only the impact of random cell-to-cell diversity in the GC population, but also the benefits of the structured heterogeneity introduced by neurogenesis.

Despite the fact that single mature GCs achieve better decoding performance than immature ones, immature GCs are selected when we build populations that optimize stimulus reconstruction. Moreover, these mixed-age populations yield better stimulus reconstruction than populations consisting of exclusively mature GCs. Correspondingly, a previous study involving spatial exploration has shown that immature GCs are less spatially tuned but actively participate in context encoding and discrimination [10]. Additionally, they reported that immature GCs fire at higher rates in vivo. In this study, we equalize the firing rates of different neurons, yet a higher firing rate would further improve the decoding performance of immature GCs and increase their importance in population coding.

We explore the capability of populations of GCs to perform pattern separation in the time domain, formulating the discrimination of time-varying signals on the scale of milliseconds. Recently, a study using perforant path stimulation studied pattern separation in different development-born hippocampal neurons but they only used pulsed stimulation [38]. Despite immature GCs variable responses, we find that mixed-age populations are able to discriminate between highly correlated stimuli better than exclusively mature populations, even in the presence of uncorrelated noise in the population. The noisier output of immature GCs that we observe could also help disrupt established memories, consistent with the hypothesis that neurogenesis induces forgetting of existing memories, facilitating the formation of new conflicting ones [39]. Computational studies of the dentate gyrus that incorporate neurogenesis have usually focused on learning aspects [40, 41]. Using networks that continually incorporate new GCs to encode novel information these studies propose that neurogenesis could prevent interference between new and old memories and aiding their discrimination.

It is intriguing that, while immature cells are less reliable individually, they still contribute to decoding in a population and improve performance in a pattern separation task. We explored the properties of selected individual GCs and found that many immature GCs that were highly selected in the populations did not perform well individually in decoding the stimulus. Thus, this rules out simple explanations for these observations and points to non-trivial synergistic effects in the mixed population. It is possible that immature GCs code aspects of the stimulus that are not represented in mature cells. This is an interesting open question for future work.

Our work adds evidence in favor of the idea that intrinsic heterogeneity is beneficial for population coding [7, 8]. Particularly, our study indicates that the intrinsic diversity immature GCs contribute to the dentate gyrus could be, by itself, beneficial for stimulus representation and discrimination. A general mechanistic understanding of how the dentate gyrus leverages immature GCs to process incoming activity is missing. Our approach could be used and combined with other experimental procedures to guide future experiments and further advance this understanding.

## Acknowledgements

We thank Emilio Kropff for valuable comments on the manuscript, and members of the Marin Burgin and Morelli labs for fruitful discussions. This work was supported by IDRC 108878 and ANPCyT grants PICT 2015 0634 and PICT 2018 0880 awarded to AMB, ANPCyT grants PICT 2017 3753 and PICT 2019 0445 awarded to LGM, and FOCEM-Mercosur (COF 03/11) awarded to IBioBA.

## Methods

### Mice and slice preparation

Ascl1-CreERT2 mice [26] were crossed with CAG-floxStop-tdTomato mice to generate Ascl1-CreERT2-Tom mice. Mice were housed with a running wheel which is known to enhance adult hippocampal neurogenesis [42]. Adult mice of either sex were injected with tamoxifen at 6-8 weeks of age. Tamoxifen was delivered intraperitoneally in two 120*µ*g per mouse gram injections in two consecutive days to achieve indelible expression of Tom in newborn GCs. Mice were anesthetized and decapitated at 4 or 5 weeks after injection, depending on the desired age of the GCs. Experimental protocol (2020-03-NE) was evaluated by the Institutional Animal Care and Use Committee of the IBioBA-CONICET according to the Principles for Biomedical Research involving animals of the Council for International Organizations for Medical Sciences and provisions stated in the Guide for the Care and Use of Laboratory Animals. Brains were removed and placed into a chilled solution containing (mM) 110 choline cloride, 2.5 KCl, 2.0 NaH_2_PO_4_, 25.0 NaHCO_3_, 0.5 CaCl_2_, 7 MgCl_2_, 20 dextrose, 1.3 sodium ascorbate, 0.6 sodium pyruvate. Acute slices 400*µ*m thick from either hemisphere were cut transversally to the longitudinal axis in a vibratome. Hippocampal slices were then transferred to a chamber containing artificial cerebrospinal fluid (ACSF; mM): 125 NaCl, 2.5 KCl, 2.3 NaH2PO4, 25 NaHCO3, 2 CaCl2, MgCl2, 1.3 Na+-ascorbate, 3.1 Na+-pyruvate, and 10 dextrose (315 mOsm). Slices were bubbled with 95% O2/5% CO2 and maintained at 30^◦^C for *>*1hr before experiments started.

### Electrophysiological Recordings

Recorded immature neurons were visually identified by fluorescence. GCs used between 28 and 30 days post injection were labeled as 4w and GCs used between 34 and 36 days post injection were labeled as 5w. The mature population encompassed unlabeled neurons localized in the outer third of the granule cell layer [15, 26]. Whole-cell current-clamp recordings were performed using microelectrodes (4–6 MΩ) filled with a potassium gluconate internal solution (in mM): 120 potassium gluconate, 4 MgCl2, 10 HEPES buffer, 0.1 EGTA, 5NaCl, 20KCl, 4ATP-tris, 0.3 GTP-tris, and 10 phosphocreatine (pH = 7.3; 290 mOsm). Recordings were obtained using Multiclamp 700B amplifiers (Molecular Devices), digitized using Digidata 1550 (Axon instruments), and acquired at 10 kHz onto a personal computer using the pClamp10 software (Molecular Devices). Before starting every protocol of stimulation, the resting membrane potential was kept at around -70 mV by passing a holding current. Passive properties were measured by injecting a small hyperpolarizing current step. Current step amplitude was adapted to each neuron and resulted in a negative membrane potential deviation of 1-4 mV.

### Fluctuating stimulus

We generated fluctuating stimuli from an Ornstein-Uhlenbeck process [43]

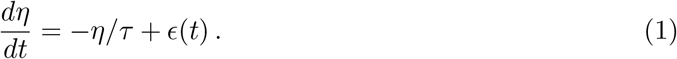

The process has zero mean, unit variance and correlation time constant *τ*. This process was implemented numerically with the discretization

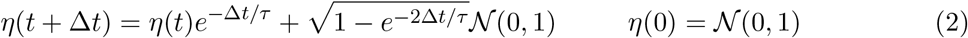

Throughout this work we used a time constant *τ* = 3 ms that was much shorter than neuron membrane time constants. Both in experiments and simulations described below, neurons were injected with currents of the form *i*(*t*) = *ση*(*t*)+*µ*, where *µ* set the mean value of the current and *σ* determined the amplitude of current fluctuations. Since GCs of different ages have markedly different input resistances, during experiments we adapted *µ* and *σ* online for each individual GC, seeking to obtain a firing rate of at least 1Hz.

### Coincidence ratio

A spike train *s*(*t*) with spike times {*t*_1_, *t*_2_, …*t*_*s*_, …} can be represented as the sum of Dirac delta distributions

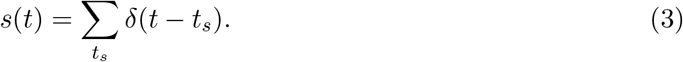

For two spike trains *s* and *s*′, we defined a coincidence between them as the co-occurrence of two spikes within a predefined time window. We introduced the number of coincident spikes *c*(*s, s*′), defined by the inner product ⟨*s, s*′⟩ between the spike trains [29, 30],

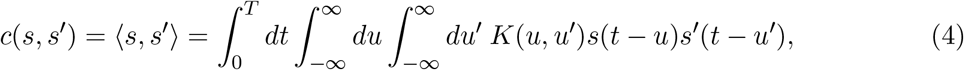

where *T* is spike train duration and *K*(*u, u*^′^) = *h*_Δ_(*u*)*δ*(*u*^′^) is a kernel with *h*_Δ_(*u*) = Θ(*u* + Δ*/*2)Θ(*u* − Δ*/*2) a rectangular function of width Δ, Θ the Heaviside step function and *δ* the Dirac delta distribution. To compute all the quantities introduced here we always used the time window Δ = 8 ms that is similar to the duration of an action potential. For this choice of kernel and the small time window used, the coincidence of a spike train *s* with itself always counted the total number of spikes *n*(*s*) in the train, ⟨*s, s*⟩ = *n*(*s*) for all spike trains considered.

Given two different sets of M spike trains *S* = {*s*^1^, *s*^2^, …, *s*^*M*^ } and *S* = {*s*^′1^, *s*^′2^, …, *s*^′*M*^ }, we define the coincidence ratio between them

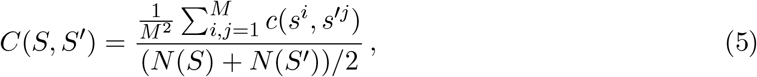

where the numerator is the average number of coincidences between the two sets and 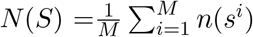 is the average number of spikes in the trains of the set *S*. This coincidence ratio is the average number of coincidences between the sets normalized by the average number of spikes in their spike trains. For a single recording *S* consisting of *M* trials, we define the reliability as

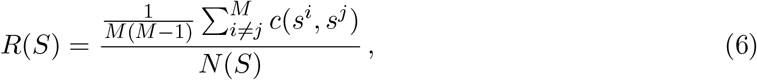

the average number of coincidences between different spike trains in the recording normalized by the average number of spikes in its spike trains. Thus, this reliability quantifies the variability within a single set of spike trains.

### Spike response model

The spike response model generates spikes from a Bernoulli probability distribution with an instantaneous rate that is determined from the subthreshold membrane potential. The sub-threshold membrane potential is

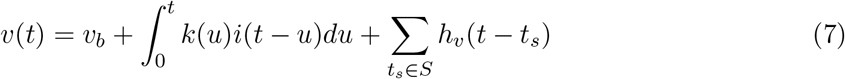

where *v*_*b*_ is a voltage bias and *k*(*t*) is a membrane filter that is convolved with the stimulus *i*(*t*). A voltage history filter *h*_*v*_(*t*) is added after every spike for the set of spike times *S*. The instantaneous rate or conditional intensity *λ*(*t*) is defined from the subthreshold membrane potential as

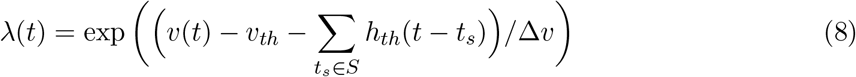

where *v*_*th*_ is a fixed voltage threshold, *h*_*th*_(*t*) is the threshold deflection every time there is a spike and Δ*v* is a voltage scale. Finally, a spike is produced at time *t* with probability

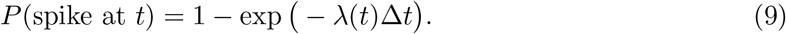

with a discretized time interval Δ*t*.

### Spike response model Fitting

To fit the spike response model (SRM) we expand the three filters *k*(*t*), *h*_*v*_(*t*) and *h*_*th*_(*t*) as linear combinations

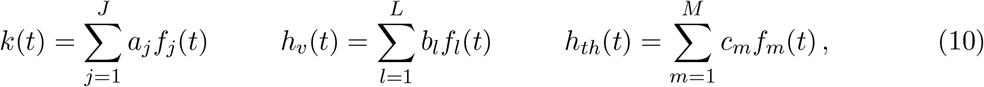

where the *f*_*j*_(*t*) are rectangular basis functions with support [*t*_*j*_, *t*_*j*+1_), that is *f*_*j*_(*t*) = 1 if *t*_*j*_ ≤ *t < t*_*j*+1_ and *f*_*j*_(*t*) = 0 otherwise. We used the same bin size for all the rectangular functions of the same filter. For *k*(*t*) we used 44 bins of 8 ms width from *t*_1_ = 0 ms to *t*_44_ = 344 ms, for *h*_*v*_(*t*) we used 17 bins of 25 ms width from *t*_1_ = 25 ms to *t*_17_ = 425 ms and for *h*_*th*_(*t*) we used 18 bins of 25 ms width from *t*_1_ = 0 ms to *t*_18_ = 425 ms. The predicted subthreshold voltage 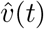 is then a linear combination of the parameters

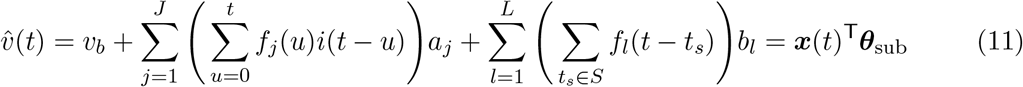

where ***x***(*t*) ∈ ℝ^1+*J* +*L*^ is the vector

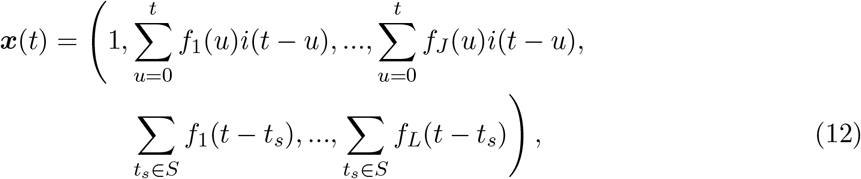

***x***(*t*)^T^ denotes the transposed vector, and ***θ***_sub_ = (*v*_*b*_, *a*_1_, …, *a*_*J*_, *b*_1_, …, *b*_*L*_) is a vector containing the subthreshold model parameters. Introducing the matrix ***X*** ∈ ℝ^*N* ×1+*J* +*L*^ of rows ***x***(*t*), we can write

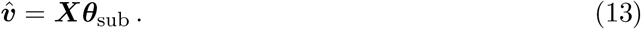

Here we distinguish the predicted 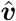 and recorded ***v*** subthreshold membrane potentials, and introduce vectors to write in compact form the time dependence at discretized times, that is ***v*** = (*v*(0), *v*(Δ*t*), …, *v*(*T*)) with *T* = (*N* − 1)Δ*t*. We determined ***θ***_sub_ from the recorded potential by using linear least squares to minimize the Mean Squared Error (MSE),

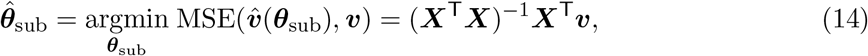

after removing the first 25 ms of voltage after every spike.

A set of spike times *S* defines a spike train ***s*** = (*s*(0), *s*(Δ*t*), …, *s*(*T*)) with *s*(*t*) = 1 if there is a spike at time *t* and *s*(*t*) = 0 otherwise. The joint probability to observe a spike train ***s*** as a function of the threshold parameters ***θ***_th_ given the stimulus ***i*** is the likelihood *P* (***s*** | ***i***; ***θ***_th_). The log-likelihood function is [30]

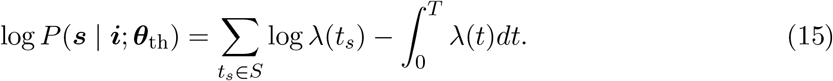

Defining

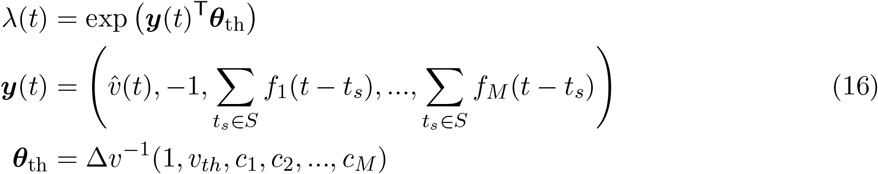

With 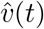 determined by Eq. (13), the log-likelihood can be written as

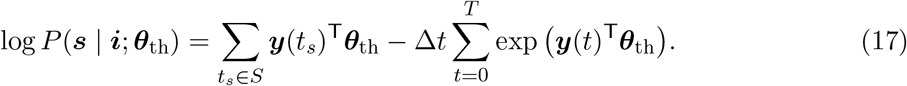

This log-likelihood is concave on the parameters and we can find its global maximum using Newton’s method to perform gradient ascent [30]. We also added a term of the form

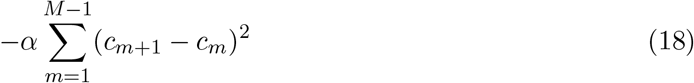

to impose a degree of smoothness determined by *α* over the coefficients of the filter *h*_*th*_(*t*). We used the value of *α* that yielded the largest log-likelihood on the validation data. We evaluated our fits by computing the root mean squared error between the predicted and recorded subthreshold voltage, the log-likelihood and a coefficient *M*_*d*_ which quantifies the degree of similarity between SRM generated and recorded sets of spike trains [29, 32]. We reported the log-likelihood relative to the log-likelihood of a Poisson process of the same rate and normalized by the number of spikes,

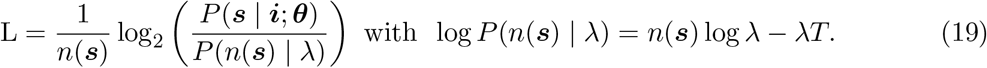

The degree of similarity *M*_*d*_ was computed using the coincidences with rectangular kernels,

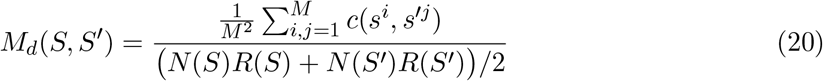

where *c*(*s, s*^′^) is given by Eq. (4), *R*(*S*) by Eq. (6) and *N* (*S*) is the average number of spikes in the trains of the set *S*. In contrast to the coincidence ratio, *M*_*d*_ takes into account the reliability within each set of spike trains.

### Decoding

Here we describe how we decoded a single stimulus ***η*** from the responses of multiple neurons. Given spike trains ***s***^*j*^ from GCs with SRM parameters ***θ***^*j*^ produced by stimuli ***i***^*j*^ = *σ*^*j*^***η*** + *µ*^*j*^, we found the maximum a posteriori estimate of ***η*** as

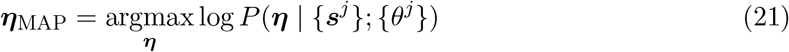

with

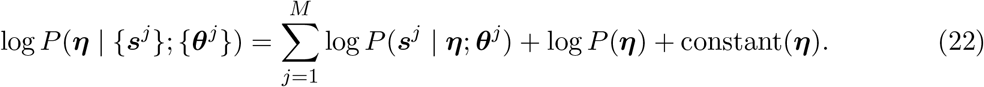

We generated stimuli using Eq. (2), resulting in

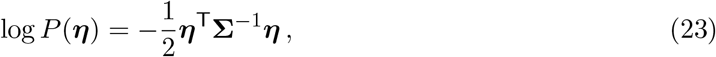

with

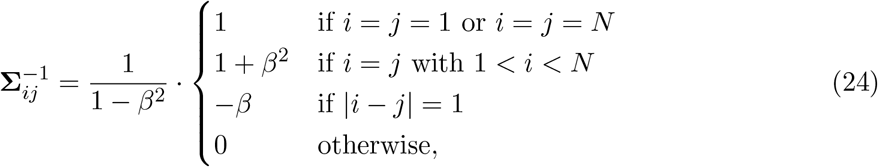

and *β* = *e*^−Δ*t/τ*^. The log-posterior of Eq. (22) with the Gaussian prior of Eq. (23) is concave on ***η***, so we found its global maximum using Newton’s method to perform gradient ascent [34]. Once we found ***η***_MAP_, approximating the posterior by a Gaussian, *P* (***η*** | {***s***^*j*^}; {***θ***^*j*^}) ≈ 𝒩 (***η***_MAP_, ***C***) we used

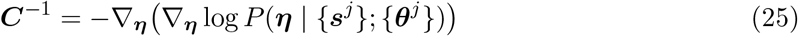

to determine the covariance matrix of the approximation and quantify uncertainty [34]. We used the squared root of the diagonal of ***C*** to quantify the uncertainty in the estimation as a function of time and the determinant of ***C*** to compute the mutual information. The matrix ***C***^−1^ is positive semi-definite, symmetric and banded. We used these properties and implemented the Cholesky decomposition from SciPy [44] to compute the optimization updates, the diagonal of ***C*** and the determinant of ***C*** efficiently and saving memory.

Given a set of GCs indexed by *j*, we sampled multiple stimuli from Eq. (2). For each stimulus we used the currents ***i***^*j*^ = *σ*^*j*^***η*** + *µ*^*j*^ to generate spike trains and performed the decoding. We quantified the reconstruction error using the coefficient of determination,

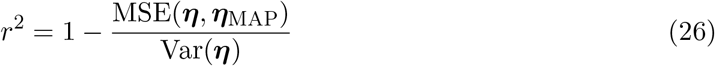

averaged over all sampled ***η***. We obtained the mutual information between the spike trains {***s***^*j*^} and the stimulus ***η***,

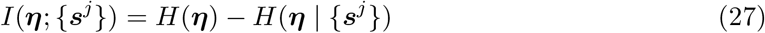

using all the decoded stimuli by computing

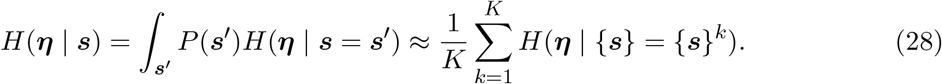

We used the Gaussian prior and Gaussian approximation of the posterior to compute the Gaussian entropies

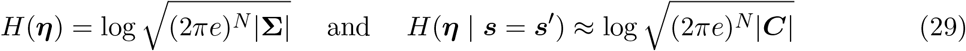

where | · | is the matrix determinant.

Where we sought to compensate for differences in firing rates, we used a different number of trials for each GC to equalize the number of spikes that each one contributes to the decoding. To obtain *n* spikes on average from a GC with firing rate *λ* in a simulation of duration *T*, we used round(*n/*(*λT*)) trials.

### Pattern discrimination

We generated pairs of correlated stimuli ***η***(*t*) = (*η*_1_(*t*), *η*_2_(*t*)) following

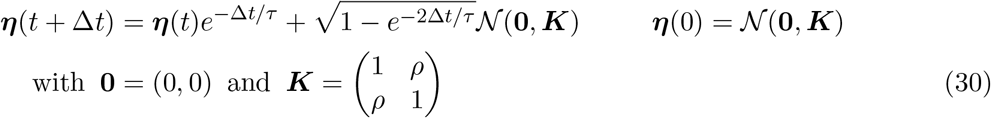

with 0 ≤ *ρ <* 1. The two processes have mean value 0, variance 1, time constant *τ* = 3 ms and correlation ⟨*η*_1_(*t*)*η*_2_(*t*)⟩ = *ρ*. We used only *η*_1_(*t*) to generate spike trains and perform the decoding. We computed the mean squared error (MSE) between the reconstruction and both stimuli *η*_1_(*t*) and *η*_2_(*t*). If MSE(***η***_MAP_, ***η***_1_) *<* MSE(***η***_MAP_, ***η***_2_) then the patterns were correctly discriminated and incorrectly otherwise. By sampling multiple pairs ***η***_1_, ***η***_2_ we quantified the accuracy of the discrimination task as the percentage of correctly discriminated stimuli. For quantitation, we introduce three degrees of separation, defined as low (*ρ* = 0.99), medium (*ρ* = 0.999) and high (*ρ* = 0.9997).

## Supplementary material

**Figure 1-figure supplement 1:**
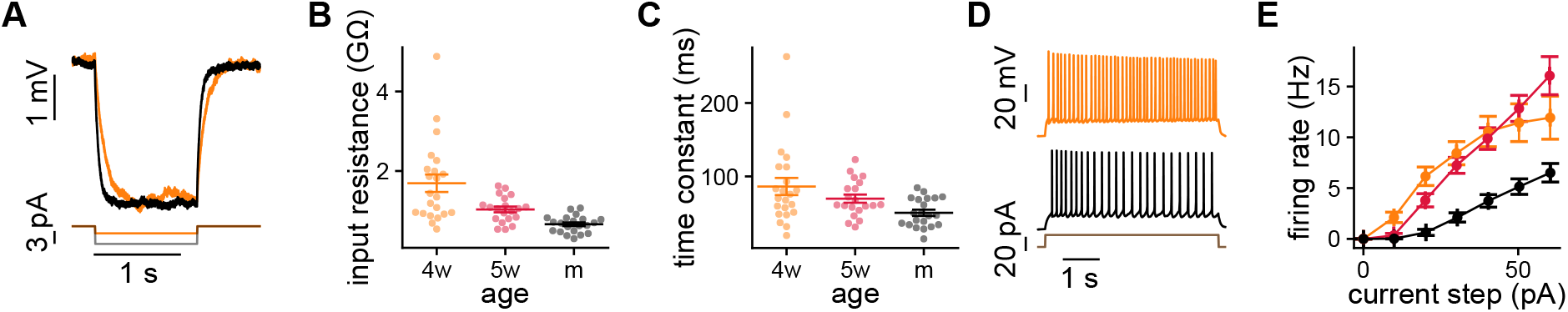
Intrinsic properties of GCs measured with current steps. **(A)** Negative step in a 4wGC (orange) and a mGC (black). **(B, C)** Passive properties obtained with negative steps. Spearman’s correlation between age and input resistance: *ρ* = −0.64, *p* = 1.4×10^−8^; Spearman’s correlation between age and time constant: *ρ* = −0.36, *p* = 4.3×10^−3^. **(D)** Positive step in a 4wGC (orange) and a mGC (black). **(E)** Firing rate vs. amplitude of the step for the different age groups.

**Figure 1-figure supplement 2:**
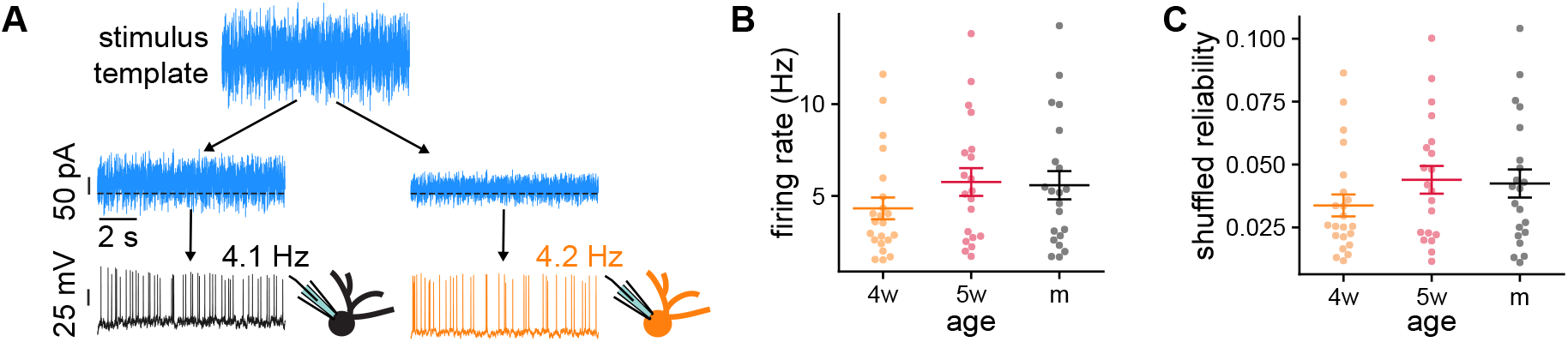
Adjusting the baseline and amplitude of the stimulus to GCs of different ages while keeping the same time structure. **(A)** Immature GCs have larger input resistances, hence they need smaller currents to produce similar firing rates. **(B)** Resulting firing rates for the recorded GCs. There was no significant difference between the groups. Kruskal–Wallis H test: *p* = 0.29. **(C)** Reliabilities obtained after shuffling the spike times. Spearman’s correlation between reliability and age was reduced in the shuffled data (*p* = 0.014). *p* was calculated by boostrapping GCs of all ages, computing Spearman’s correlation with age of the reliability and the shuffled reliability and taking the fraction of realizations in which the correlations were larger in the shuffled data.

**Figure 2-figure supplement 1:**
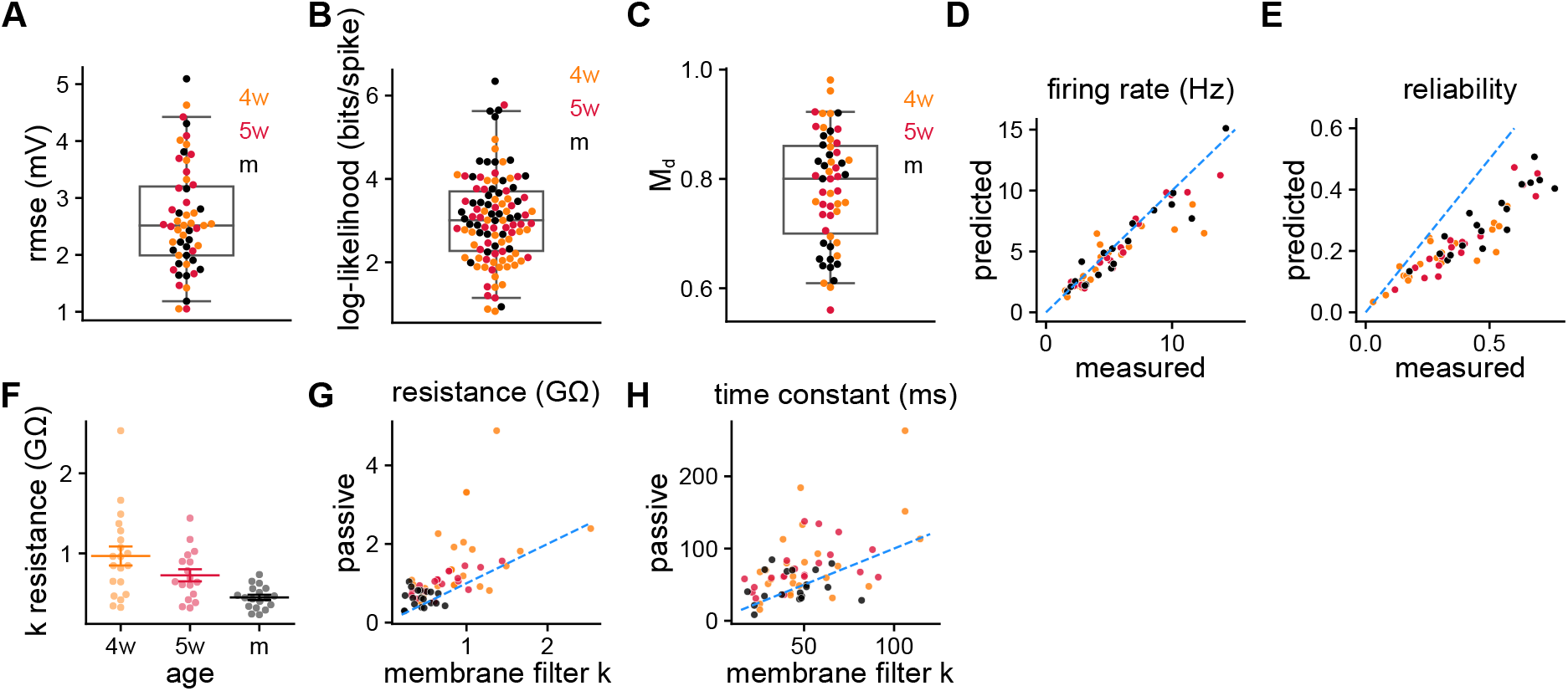
**(A)** Root mean squared error of the subthreshold membrane potential prediction using the validation data. **(B)** Log-likelihood per spike relative to the likelihood of a Poisson process of the same rate measured on the validation data. **(C)** *M*_*d*_ between recorded validation data and model-generated spike trains. **(D)** Firing rate and **(E)** reliability recorded and obtained by generating spike trains with the SRM. Pearson’s correlation between predicted and measured reliability: *ρ* = 0.92, *p* = 7 × 10^−24^. **(F)** Electrical resistance obtained from the membrane filter *k* (Figure 2C). Spearman’s correlation between age and *k* resistance: *ρ* = −0.52, *p* = 1.0 × 10^−4^. **(G)** Passive resistance and **(H)** time constant vs. the ones obtained from the membrane filter *k*(*t*). Wilcoxon signed-rank test between passive and *k* resistance: *p* = 8.6×10^−9^; Spearman’s correlation between age and *k* resistance after subtracting age groups: *ρ* = 0.52, *p* = 1.1 × 10^−5^. Wilcoxon signed-rank test between passive and *k* time constant: *p* = 3.5 × 10^−5^.

**Figure 2-figure supplement 2:**
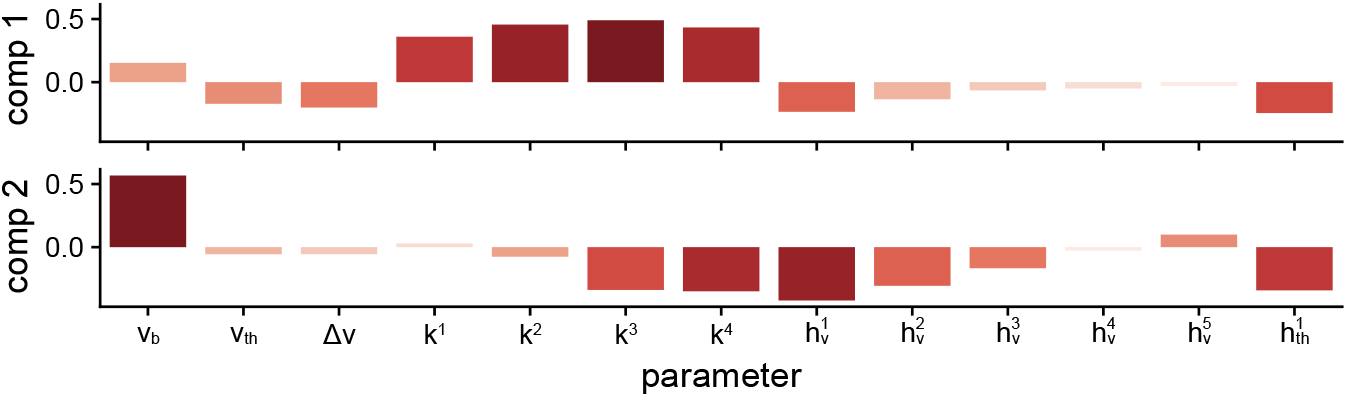
Linear Discriminant Analysis components as determined by the scalings of each parameter used.

**Figure 3-figure supplement 1:**
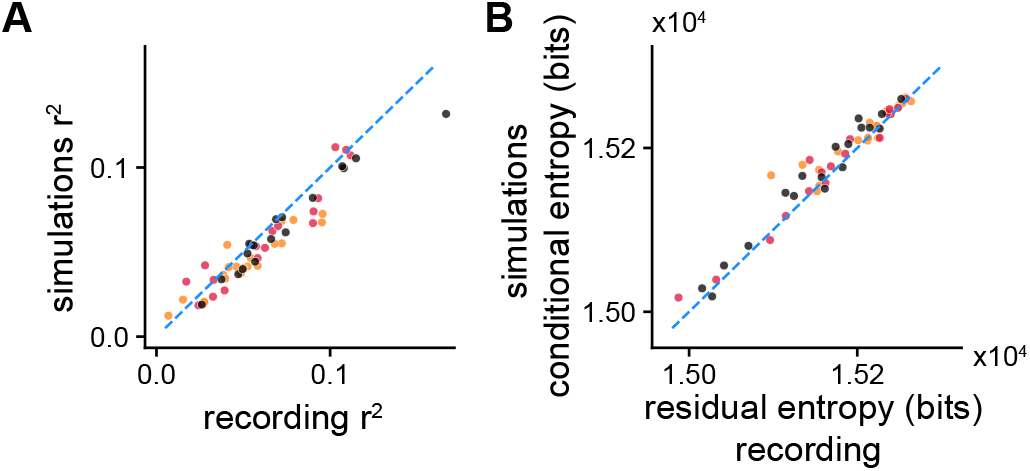
**(A)** *r*^2^ and **(B)** information from Figures 3D,E vs. the *r*^2^ and reduction in uncertainty obtained by decoding the experimentally used stimulus from the recorded spike trains.

**Figure 3-figure supplement 2:**
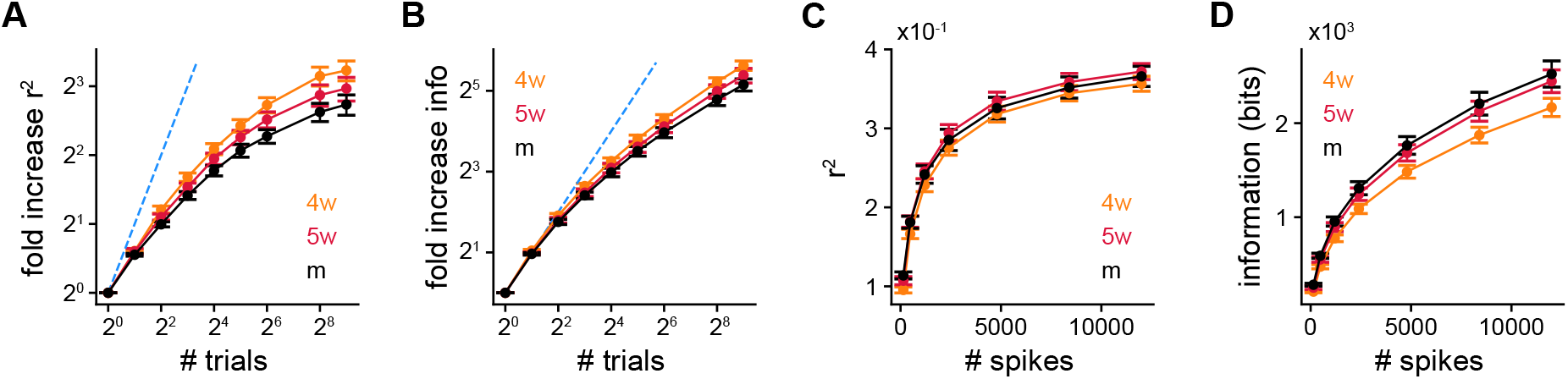
Fold increase in **(A)** *r*^2^ and **(B)** information with number of trials from single GCs. Spearman’s correlation between age and *r*^2^ at 2^6^ trials: *ρ* = −0.42, *p* = 1.5 × 10^−3^; Spearman’s correlation between age and information at 2^6^ trials: *ρ* = −0.36, *p* = 6.3 × 10^−3^. **(C)** *r*^2^ and **(D)** information using multiple trials from single GCs while using on average the same number of spikes from each one of them. Kruskal-Wallis H test between age and *r*^2^ for all number of spikes: *p >* 0.27; Spearman’s correlation between age and information at 140 spikes: *ρ* = 0.44, *p* = 8.5 × 10^−4^; Spearman’s correlation between age and information at 12000 spikes: *ρ* = 0.26, *p* = 5.1 × 10^−2^.

**Figure 3-figure supplement 3:**
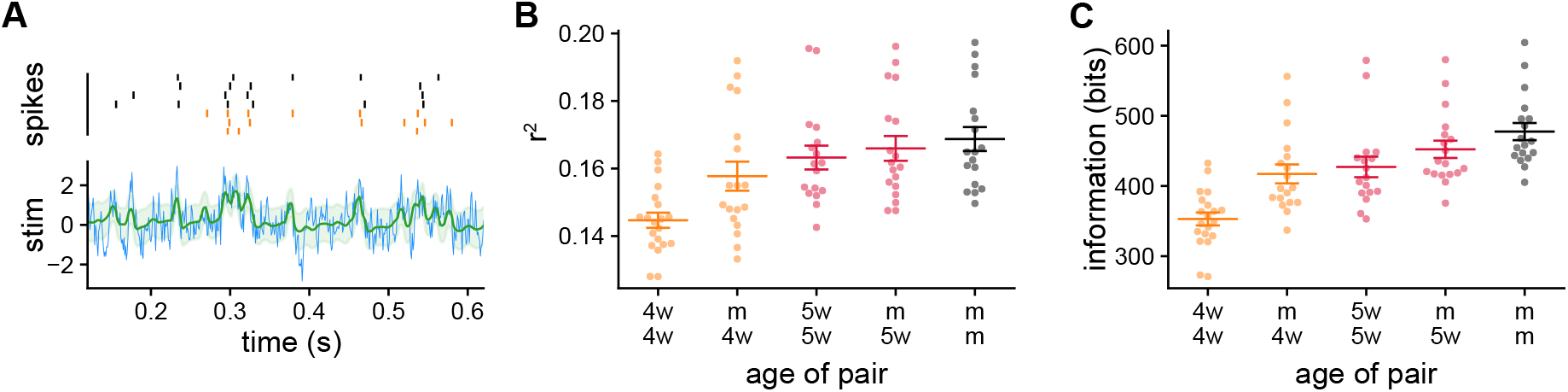
**(A)** Decoding example using approximately 140 spikes from a 5wGC and a mGC to get a single stimulus reconstruction. **(B)** *r*^2^ obtained by using pairs of GCs to decode. Spearman’s correlation between same age pair and *r*^2^: *ρ* = 0.46, *p* = 3.6 × 10^−4^; Spearman’s correlation for pairs with a mature GC between age of the second GC and *r*^2^: *ρ* = 0.21, *p* = 0.14. **(C)** Information obtained by using pairs of GCs to decode. Spearman’s correlation between same age pair and information: *ρ* = 0.75, *p* = 5.5 × 10^−11^; Spearman’s correlation for pairs with a mature GC between age of the second GC and information: *ρ* = 0.42, *p* = 1.5 × 10^−3^.

**Figure 4-figure supplement 1:**
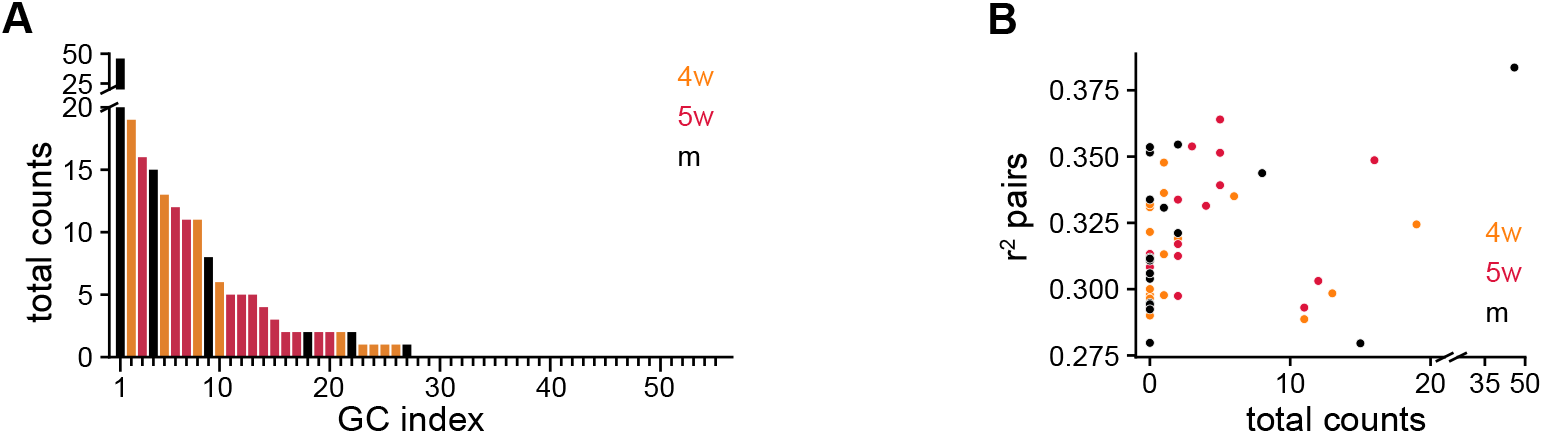
**A)** Total number of times each GC was selected by the greedy procedure after 12 steps. **(B)** Average *r*^2^ value achieved by each GC when paired with every other possible GC vs. total number of times the GC was selected. Each GC in the pair used approximately 1200 spikes.

